# Uncovering the gene machinery of the Amazon River microbiome to degrade rainforest organic matter

**DOI:** 10.1101/585562

**Authors:** Célio Dias Santos, Hugo Sarmento, Fernando Pellon de Miranda, Flávio Henrique-Silva, Ramiro Logares

**Author notes:** **Corresponding authors:** FHS, RL.

## Abstract

The Amazon River receives, from the surrounding rainforest, huge amounts of terrestrial organic matter (TeOM), which is typically resistant to microbial degradation. However, only a small fraction of the TeOM ends up in the ocean, indicating that most of it is degraded in the river. So far, the nature of the genes involved in TeOM degradation and their spatial distributions are barely known. Here, we examined the Amazon River microbiome gene repertoire and found that it contains a substantial gene-novelty, compared to other environments (rivers and rainforest soil). We predicted ~3.7 million non-redundant genes, affiliating mostly to bacteria. The gene-functions involved in TeOM degradation revealed that lignin degradation correlated to tricarboxylates and hemicellulose processing, pointing to higher lignin degradation rates under consumption of labile compounds. We describe the biochemical machinery that could be speeding up the decomposition of recalcitrant compounds in Amazonian waters, previously reported only in incubation experiments.

## INTRODUCTION

Continental waters play a major biogeochemical role by linking terrestrial and marine ecosystems^1^. Riverine ecosystems receive terrestrial organic carbon, which is mostly processed by microorganisms, stimulating the conversion of terrestrially derived organic matter (TeOM), which can be recalcitrant, to carbon dioxide^2–4^. Therefore, riverine microbiomes should have evolved metabolisms capable of degrading TeOM. Even though the gene repertoire of river microbiomes can provide crucial insights to understand the links between terrestrial and marine ecosystems, as well as the fate of organic matter synthesized on land, very little is known about the genomic machinery of riverine microbes that degrade TeOM.

Microbiome gene catalogues allow the characterization of the functional repertoire, linking genes with ecological function and ecosystem services. Recently, large gene catalogues have been produced for the global ocean^5–7^, soils^8^ and animal guts^9, 10^. In particular, ~40 million genes have been reported for the global ocean microbiome^7^ and ~160 million genes for the global topsoil microbiome^8^.

So far, there is no comprehensive gene catalogue for rivers, which hinders our comprehension of the genomic machinery that degrade almost half of the 1.9 Pg C of recalcitrant TeOM that are discharged into rivers every year^1^. This is particularly relevant in tropical rainforests, like the Amazon forest, which accounts for ~10% of the global primary production, fixating 8.5 Pg C per year ^11, 12^. The Amazon River basin comprises almost 38% of continental South America^13^ and its discharge accounts for 18% of the world’s inland-water inputs to the oceans^14^. Despite its relevance for global scale processes, there is a limited understanding of the Amazon River microbiome, as well as microbiomes from other large tropical rivers.

The large amounts of organic and inorganic particulate material^15^ turns the Amazon River into a turbid system. High turbidity reduces light penetration and, consequently, the Amazon River has very low rates of algal production^16^, meaning that the dissolved organic carbon cycling at the terrestrial–aquatic interface is the major carbon source for microbial growth^17^. High respiration rates in Amazon River waters generate a super-saturation that leads to CO_2_ outgassing to the atmosphere. Overall, Amazon River outgassing accounts for 0.5 Pg C per year to the atmosphere^18^, almost equivalent to the amount of carbon sequestered by the forest^11, 12^. Despite the predominantly recalcitrant nature of the TeOM that is discharged into the Amazon River, heterotrophic microbes are able to degrade up to 55% of the lignin produced by the rainforest^19, 20^. The unexpectedly high degradation rates of some TeOM compounds in the river was recently explained by the availability of labile compounds that promote the degradation of recalcitrant ones, a mechanism known as *priming effect*, which has been observed in incubation experiments^20^.

Determining the repertoire of gene-functions in the Amazon River microbiome is one of the key steps to understand the mechanisms involved in the degradation of complex TeOM produced by the rainforest. Given that most TeOM present in the Amazon River is lignin and cellulose^19–23^, the functions associated to their degradation were expected to be widespread in the Amazon microbiome. Instead, these functions exhibited very low abundances^24–26^, highlighting our limited understanding of the enzymes involved in the degradation of lignin and cellulose in aquatic systems.

Cellulolytic bacteria use an arsenal of enzymes with synergistic and complementary activities to degrade cellulose. For example, glycosyl-hydrolases (GHs) catalyze the hydrolysis of glycoside linkages, polysaccharide esterases support the action of GHs over hemicelluloses, and polysaccharide lyases promote depolymerisation^27, 28^. In contrast, lignin is more resistant to degradation^29^, since its role is preventing microbial enzymes from degrading labile cell-wall polysaccharides^30^. The microbial production of extracellular hydrogen-peroxide, a highly reactive compound, is the first step of lignin oxidation mediated by enzymes, like lignin peroxidase, manganese-dependent peroxidase and copper-dependent laccases^31^. Lignin oxidation also produces a complex mixture of aromatic compounds, which compose the humic fraction of dissolved carbon detected in previous studies in the Amazon River mainstream^21, 22^. Lignin degradation tends to occur in oxic waters of the Amazon River, using the hydrogen peroxide produced by the metabolism of cellulose and hemicellulose^32^. Therefore, a higher amount of lignin degradation genes is expected in oxic waters.

Here, we produced the first gene catalogue of the world’s largest rainforest river by analyzing 106 metagenomes (~500 ×10^9^ base pairs), originating from 30 stations covering a total of ~ 2,106 km, from the upper Solimões River to the Amazon River plume in the Atlantic Ocean. This gene catalogue was used to uncover and examine the genomic machinery of the Amazon River microbiome to metabolize large amounts of carbon originating from the surrounding rainforest. Specifically, we ask: How novel is the gene repertoire of the Amazon River microbiome? Which are the main functions associated to TeOM degradation? Do TeOM degradation genes and functions have a spatial distribution pattern? Is there any evidence of priming effect in TeOM degradation?

## RESULTS

### Cataloguing the genes of the Amazon River microbiome

Our original dataset contained 106 metagenomes from 30 different stations (**Fig. 1a**) covering ~ 2,106 km of the Amazon River and its continuum over the Atlantic Ocean. Metagenome assembly yielded 2,747,383 contigs ≥ 1,000 base pairs, in a total assembly length of ~ 5.5×10^9^ base pairs (**Supplementary Table 1**). We predicted 6,074,767 genes longer than 150 bp, allowing also for alternative initiation codons. After redundancy elimination through clustering genes with an identity >95% and an overlap >90% of the shorter gene, the *Amazon river basin Microbial non-redundant Gene Catalogue* (AMnrGC) included 3,748,772 non-redundant genes, with half of the genes with a length ≥867 bp. About 52% of the AMnrGC genes were annotated with at least one database (**Fig. 1b**), while ~86% of the annotated genes were simultaneously annotated using two or more different databases. The gene and functional diversity recovered seemed to be representative of the total diversity present in the sampling sites, as indicated by the accumulation curves, which tended towards saturation (**Fig. 1c**).

**Figure 1.**
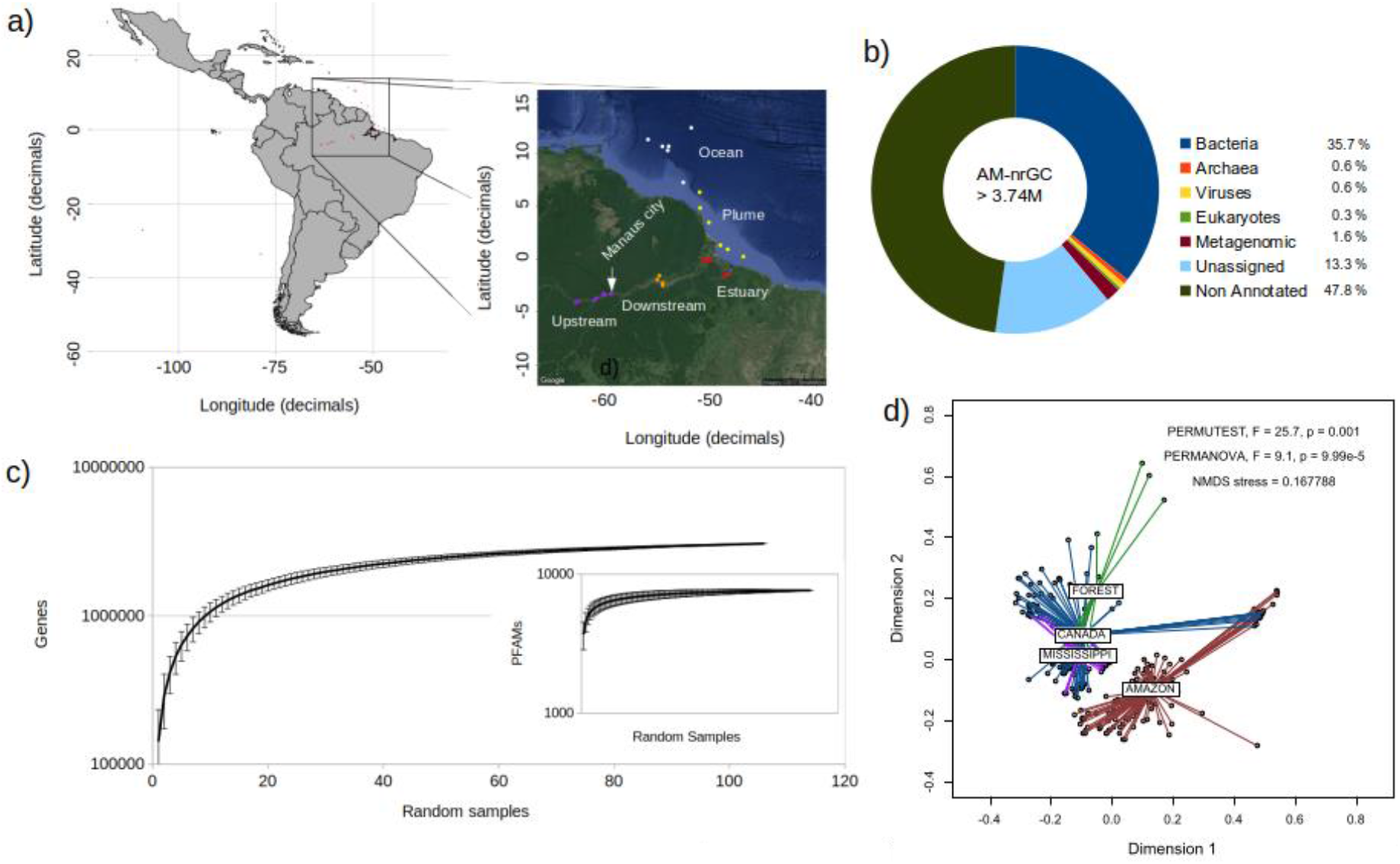
The Amazon River Basin Microbial Non-Redundant Gene Catalogue (AMnrGC). a) Distribution of the 106 metagenomes used in this work over the five sections of the Amazon River: Upstream (purple dots), Downstream (orange dots), Estuary (red dots), Plume (yellow dots) and coastal Ocean (white dots). b) Taxonomic classification of the ~ 3.7 million genes in the AMnrGC. “Unassigned” genes were not assigned taxonomy, but they were functionally assigned, differently from “non-annotated” genes, which do not have any ortholog. Those genes displaying orthology to poorly characterized genes found in metagenomes were referred as “Metagenomic”. c) Accumulation curves of non-redundant genes and PFAM families (internal graphic) point towards saturation. d) NMDS comparing the Amazon river microbiome with other microbiomes based on information content [K-mer composition; Amazon river (AMAZON), Amazon forest soil (FOREST), Canada watersheds (CANADA) and Mississippi river (MISSISSIPI)].

### Patterns in the metagenomic composition of microbiomes

We compared the metagenomic information contained in the Amazon River microbiome with that of Amazon rainforest soil and other rivers (Canada watersheds and Mississippi river). The k-mer comparison of microbiomes indicated they are different (**Fig.1d**), forming groups of heterogenous composition (significant β dispersion [that is, average distance of samples to the group centroid] - PERMUTEST, F = 25.7, p < 0.001). The metagenomic content of Amazon basin samples was different to the other compared microbiomes (PERMANOVA, R^2^ = 0.10, p = 9.99×10^−5^; ANOSIM, R = 0.27, p < 0.001), which suggests that this basin, or tropical rivers in general, contain specific gene repertoires. The metagenomic composition of the five sampled sections of the Amazon River (Upstream, Downstream, Estuary, Plume and Ocean) were significantly different (PERMANOVA test, F = 1.52, p < 9.9e-5), indicating that they do represent different assemblages from a genetic perspective. Each of these groups was considered homogenous, since there was a non-significant β dispersion (F = 2.3, p = 0.063) among metagenomic samples in each group (**Supplementary Fig. 1**).

### Gene identification

About 48% of the AMnrGC genes could not be annotated, due to lack of orthologs. Besides, even though ~1.6% of the genes in the AMnrGC were previously found in metagenomic studies, they were poorly characterized, without being assigned to a particular taxon (here referred to as “Metagenomic” genes; **Fig. 1b**). Genes annotated exclusively through Hidden Markov Models (HMM) represented 13.3% of AMnrGC (**Fig. 1b**). As the annotation using HMM profiles does not rely on direct orthology to specific sequences, but on orthology to a protein family (which may include mixed taxonomic signal), we could not assign taxonomy to those genes and they are referred as “Unassigned genes” (**Fig. 1b**).

The previous highlights our limited understanding about the gene composition of the Amazon River microbiome, where most proteins (61.11%) do not have orthologs in main reference databases. Taxonomically assigned prokaryotic genes (35.7% bacterial and 0.6% archaeal) constituted the majority in the AMnrGC, with only 0.3% and 0.6% of the genes having eukaryotic or viral origin, respectively (**Fig. 1b**).

### General or core metabolisms

The superclass “Metabolic processes” from the Clusters of Orthologous Genes (COG) database comprises those gene-functions belonging either to energy production and conversion; amino acids, nucleotides, carbohydrates, coenzymes, lipids and inorganic ions transport and metabolism; and secondary metabolites biosynthesis, transport, and catabolism. This superclass was the most abundant in the AMnrGC (35.8% of the genes annotated with COG classes), **Fig. 2**. Genes with unknown function represented 21.4% of the COG-class annotated proteins.

**Figure 2.**
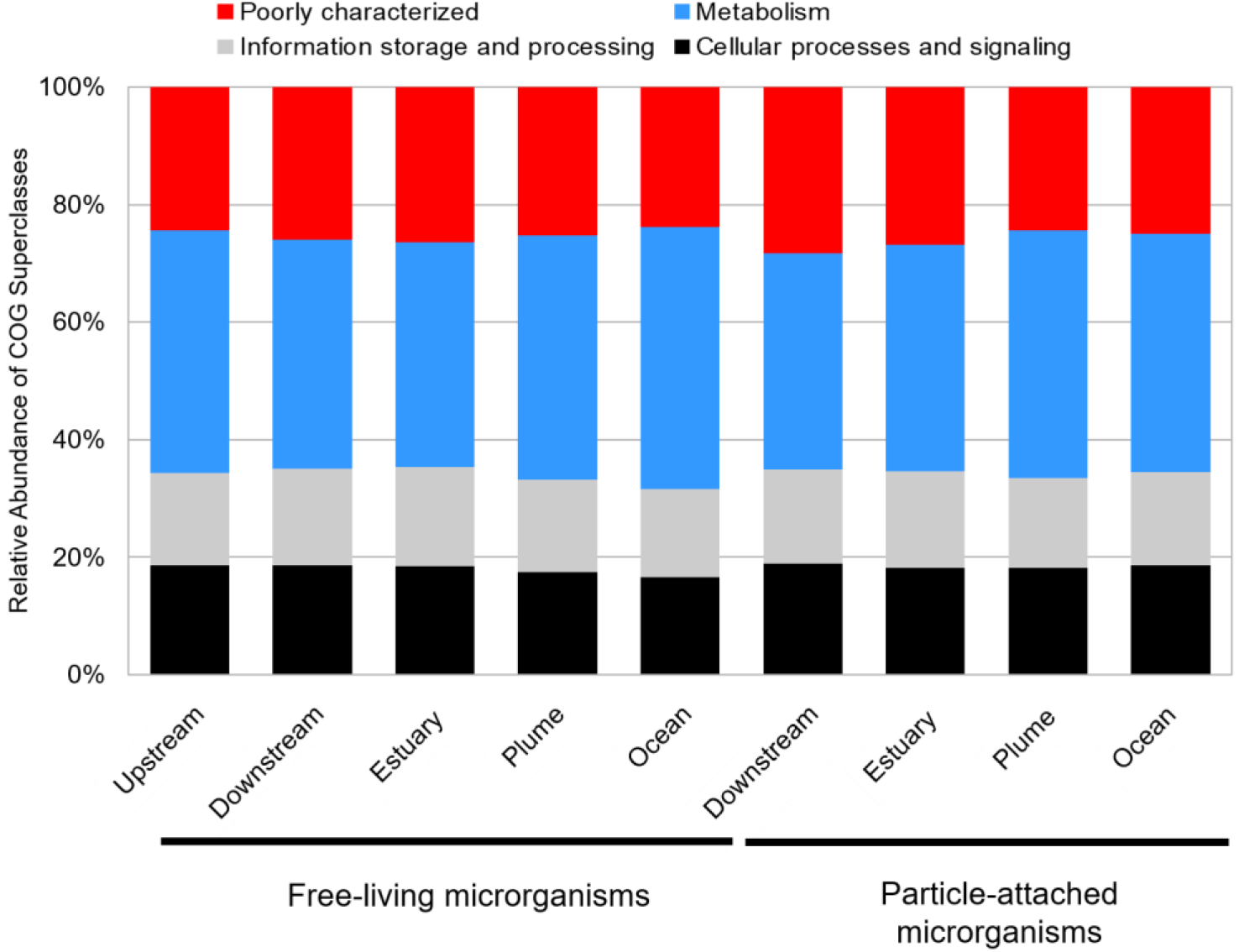
Functional composition across microbial lifestyles and sections of the Amazon River. Gene functions grouped into COG super classes are shown per river section and microbial lifestyle (particle-attached vs. free-living). Functions related to the metabolism super class were more represented in free-living that in particle-attached communities (p < 0.05, Mann-Whitney U Test). In fresh- and brackish water, all COG classes were differentially distributed, with higher gene diversities observed in freshwaters (p < 0.01, Mann-Whitney U Test). The Upstream river section is not shown in the particle-associated fraction, since it was not sampled.

Metabolism core functions were defined as those involved in cell or ecosystem homeostasis, normally representing the minimal metabolic machinery needed to survive in a given environment. KEGG and PFAM databases were used to determine the bacterial functional core, also allowing the identification of metabolic pathways. Core functions represented ~8% of KEGG and PFAM functions, and were mostly related to a general carbon metabolism, mostly associated to general organic matter oxidation until CO_2_ and the microbial respiration byproducts heading to acetogenic pathways. Besides the core, the most abundant proteins can reveal essential biochemical pathways in microbiomes. The top 100 most abundant functions in the bacterial core were “house-keeping” functions involved in main metabolic pathways (e.g. carbohydrate metabolism, *quorum sensing*, transporters and amino-acid metabolism), as well as important protein complexes (e.g. RNA and DNA polymerases and ATP synthase).

### TeOM degradation machinery

A total of 6,516 genes from the AMnrGC were identified as taking part in the TeOM degradation machinery of the Amazon River microbiome, being divided into: cellulose degradation (143 genes), hemicellulose degradation (92 genes), lignin oxidation (73 genes), lignin-derived aromatic compounds transport and metabolism (2,324 genes) and tricarboxylate transport (3,884 genes) [**Supplementary Fig. 2**]. The huge gene diversity associated to metabolism of lignin-derived compounds and the transport of tricarboxylates reflects the molecular diversity of the compounds generated, respectively, in the lignin oxidation process and present in Amazon freshwaters as humic and fulvic acids.

### Initial steps of TeOM degradation: lignin oxidation and deconstruction of cellulose and hemicellulose

TeOM consists of biopolymers, so the first step of its microbial degradation consists in converting polymers into monomers. Thus, the identified genes involved in the oxidation of lignin and degradation of cellulose and hemicellulose were investigated (**Supplementary Fig. 2**). We found that the lignin oxidation in the Amazon River is mainly mediated by dye-decolorizing peroxidases (DyPs) and predominantly associated to freshwaters. Only laccases and peroxidases were found in the Amazon River microbiome, no other families involved in lignin oxidation, like phenolic acid decarboxylase or glyoxal oxidase, were found. In turn, hemicellulose degradation seems to be performed mainly by glycosyl hydrolase GH10 in all river sections. We observed a similar ubiquitous dominance of glycosyl hydrolase GH3 in cellulose degradation across river sections. Interestingly, according to the gene content, cellulose and hemicellulose degradation seemed to replace lignin oxidation in brackish waters, suggesting the aging of TeOM during its flow through the river.

### Lignin-derived aromatic compounds degradation

Following the initial degradation of lignin, plenty of aromatic compounds are released. These can be divided into aromatic monomers (monoaryls) or dimers (diaryls), which can be processed through several biochemical steps (also called funneling pathways) until being converted into vanilate or syringate. These compounds can be processed through the O-demethylation/C1 metabolism and ring cleavage pathways to form pyruvate or oxaloacetate, which can be incorporated to the TCA cycle of the cells, generating energy. The genes identified in the AMnrGC belonging to these pathways were examined.

All known functions taking place in the metabolism of lignin-derived aromatic compounds were found in the AMnrGC, except the gene *ligD*, a Cα-dehydrogenase for αR-isomers of β-aryl ethers. The complete degradation pathway of lignin-derived compounds (**Supplementary Fig. 2d**) included 772 and 449 genes belonging to funneling pathways of diaryls and monoaryls, respectively. Examination of the pathways starting with vanilate and syringate revealed 346 genes responsible for the O-demethylation and C1-metabolism steps, while 713 genes seemed responsible for the ring-cleavage pathway. Almost 47% of all genes related to the degradation of lignin-derived compounds in the AMnrGC belonged to 4 gene families (*ligH, desV, phcD* or *phcC*). These genes represent the main steps of intracellular lignin metabolism, which are, 1) funneling pathways leading to vanilate/syringate, 2) O-demethylation/C1 metabolism and 3) ring cleavage.

We evaluated whether genes associated to TeOM degradation had a spatial distribution pattern along the river course. For this, we used the linear geographic distance of samples from the Amazon River source in Peru as a reference. Distance was negatively correlated with the number of genes associated to lignin oxidation, hemicellulose degradation, ring cleavage pathway, tripartite tricarboxylate transporting and the AAHS transporters (**Fig. 3**). This suggests a potential reduction of the microbial gene repertoire related to lignin processing as the river approaches the ocean.

**Figure 3.**
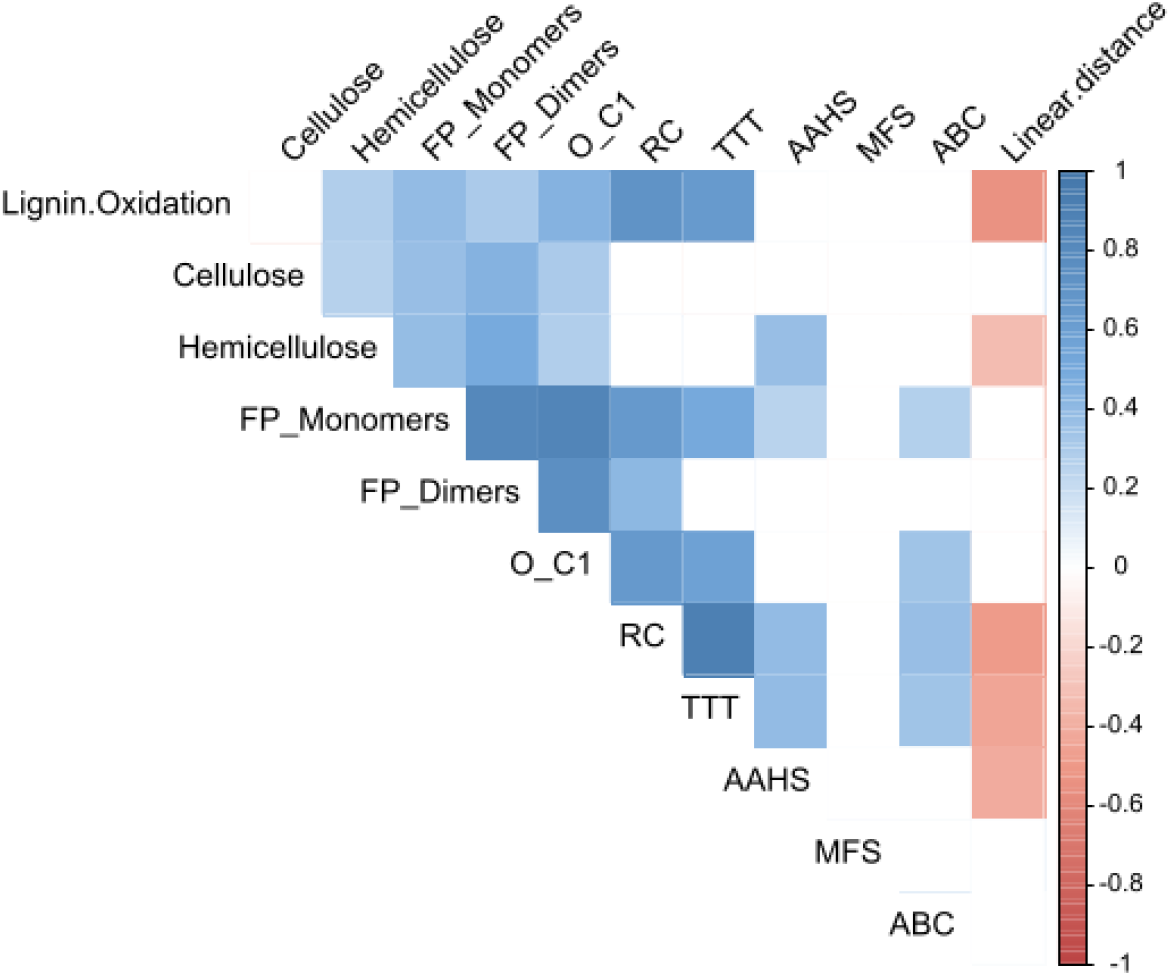
Correlations among genes associated to the processing of TeOM and geographic distance in the Amazon River. Correlations between the number of genes associated to lignin oxidation (Lignin.Oxidation), cellulose and hemicellulose deconstruction (cellulose and hemicellulose, respectively), transporting systems (AAHS, MFS, ABC and TTT), lignin-derived aromatic compounds processing pathways (RC: Ring cleavage pathways; O_C1: O demethylation / C1 metabolism pathways; Funneling pathways of Dimers - FP_Dimers and Monomers - FP_Monomers), and linear geographic distance using the river source as a starting point (Linear.distance). Color indicates correlation strength. Only significant correlations (p < 0.01) are shown.

The gene machinery associated to the processing of lignin-derived aromatic compounds was positively correlated to lignin oxidation along the river course (**Fig. 3**), suggesting a co-processing of lignin and its byproducts. Lignin oxidation and hemicellulose degradation pathways were positively correlated (**Fig. 3**), supporting the idea that monomers of hemicellulose, mainly carbohydrates, could be priming lignin oxidation. Cellulose degradation was not correlated with lignin oxidation, but had a weak positive correlation to hemicellulose degradation (**Fig. 3**), suggesting a coupling between both pathways.

We did not find correlations between the different types of funneling pathways (FP_Dimers and FP_Monomers) and the linear geographic distance along the river course (**Fig. 3**). This indicates that the degradation of lignin-derived aromatic compounds was not restricted to any river section. Moreover, the number of genes related to hemi-/cellulose degradation was positively correlated to lignin-derived aromatic compounds degradation pathways, revealing a potential co-metabolism of lignin-derived compounds and hemi-/cellulose degradation, instead of lignin-oxidation.

### Transporters

Lignin-derived aromatic compounds need to be transported from the extracellular environment to the cytoplasm prior to their degradation. Transporters that could be associated to lignin degradation (MFS transporter, AAHS family and ABC transporters) were found in the AMnrGC, while transporters from the MHS family, ITS superfamily and TRAP could not be found. MFS transporters were not correlated to any of the other examined pathways. AAHS transporters were negatively correlated to linear geographic distance, while the other transporter families did not show any type of correlation with distance (**Fig. 3**). Furthermore, AAHS and ABC transporters showed positive correlations to the funneling pathway of monoaryls, suggesting their transport by those transporter families. ABC transporters were positively correlated to O-demethylation and C1 metabolism, while AAHS and ABC transporters were correlated to the ring cleavage pathway. This suggests that ABC and AAHS transporters are relevant for the metabolism of monoaryls derived from lignin oxidation.

The tripartite tricarboxylate transporting (TTT) system was correlated to the processing of allochthonous organic material in the Amazon River. Three proteins compose this system, where one is responsible of capturing substrates in the extracellular space and bringing them to the transporting channel made by the other two proteins, which recognize the substrate binding protein and internalize the substrate. Interestingly, there is a huge diversity associated to the substrate binding proteins, since each protein is specific to one or a few substrates. Furthermore, the TTT system displayed a negative correlation with the linear geographic distance, suggesting its predominance in freshwaters sections (**Fig. 3**).

The TTT system was positively correlated to AAHS and ABC transporters (**Fig. 3**) suggesting functional complementarity, as the TTT would transport substrates not transported by the other transporter families. Furthermore, the TTT transporters showed a positive correlation with lignin oxidation and hemicellulose degradation, suggesting either the transport of the products of those processes by TTT family or a dependence of compounds transported by it.

## DISCUSSION

The AMnrGC represents the first inland tropical water non-redundant microbial gene catalogue. It allowed us to expand considerably our comprehension of the world’s largest river microbiome. Half of the ~3.7 millions genes in the AMnrGC had no orthologs, suggesting gene novelty. Yet, there is limited information about the gene repertoire in other rivers, preventing exhaustive comparisons. The analysis of k-mers indicated a distinct metagenomic composition in the Amazon River basin when compared to other rivers and to the Amazon rainforest soil. This suggests that evolutionary processes may have generated such diversity via local adaptation, although more samples from other rivers throughout the world would be necessary to test this hypothesis fully.

As expected, COG functions within the superclass “Metabolism” were the most abundant in the AMnrGC as well as in the upper Mississippi River^34^. A large fraction of these functions likely represents “core functional traits” shared across the tree of life. This was supported by the similar distribution of COGs along different sections of the Amazon River, which also points to “core functional traits” that are conserved throughout the river course. A set of core functions was also reported for the Mississippi River^34^ as well as for the global Ocean^7^.

We observed a subset of gene functions present in fresh- and brackish water sections, pointing to common core functions present along the Amazon River basin. Yet, other genes displayed a heterogeneous distribution, pointing to salinity as a structuring variable. Salinity is known to affect microbial spatiotemporal distribution, and jumps across the salinity barrier are rare evolutionary events^35^. The plume section displayed higher gene diversity than the ocean, probably reflecting the coalescence of freshwater and marine microbial communities and their different genes^36^.

Core functions included a general carbohydrate metabolism and several transporter systems, mainly ABC transporters. This suggests a sophisticated machinery to process TeOM in the Amazon River, where TeOM degradation appears more related to acetogenic and methanotrophic pathways. This agrees with previous findings^24^, indicating a high expression of C1 metabolism genes (methane monooxygenase - *mmoB* and formaldehyde activating enzyme - *fae*). The non-core pathways suggest adaptations to a complex environment, including multiple genes related to xenobiotic biodegradation and secondary metabolism (that is, the production and consumption of compounds not directly related to cell survival).

Lignin-derived aromatic compounds need to be transported from the extracellular milieu to the cytoplasm to be degraded, and different transporting systems can be involved in this process^37, 38^. In particular, previous studies showed that the TTT system was present in high quantities in the Amazon River, and this was attributed to a potential degradation of allochthonous organic matter^33^. Recent findings also suggest a TTT system related to the transport of TeOM degradation byproducts^39, 40^. Little is known about these transporters, but our findings indicate that TTT is an abundant protein family in the Amazon River, suggesting that tricarboxylates are a common carbon source for prokaryotes in these waters. Our results suggest that the TTT transporters could be linked to lignin oxidation and hemicellulose degradation, supporting their role in TeOM degradation.

Based in our findings, we propose a model of the potential priming effect acting in ligno-cellulose complexes from the Amazon River (**Fig. 4**). In this model, there are two different communities co-existing in a consortium: one responsible for hemi-/cellulose degradation and another one responsible for lignin degradation. The first community releases extracellular enzymes (mainly glucosyl hydrolases from families GH3 and GH10), whose reaction produces carbohydrates. These sugars can provide structural carbon and energy for themselves and for the lignin degrader community. The lignolytic community can also use the cellulolytic byproducts to growth, promoting an oxidative metabolism. This oxidative metabolism triggers the production and secretion of reactive oxygen species (ROS). ROS are then used by DYPs and laccases secreted by lignolytic communities to oxidize lignin, exposing more hemi-/cellulose to cellulolytic communities and re-starting the cycle. Another important role of lignolytic communities is the degradation of lignin-derived aromatic compounds generated by the lignin oxidation. If those compounds are not degraded, they could inhibit cellulolytic enzymes and microbial growth^41–44^, preventing TeOM degradation. This cycle may be considered as a priming effect, where both communities benefit from each other.

**Figure 4.**
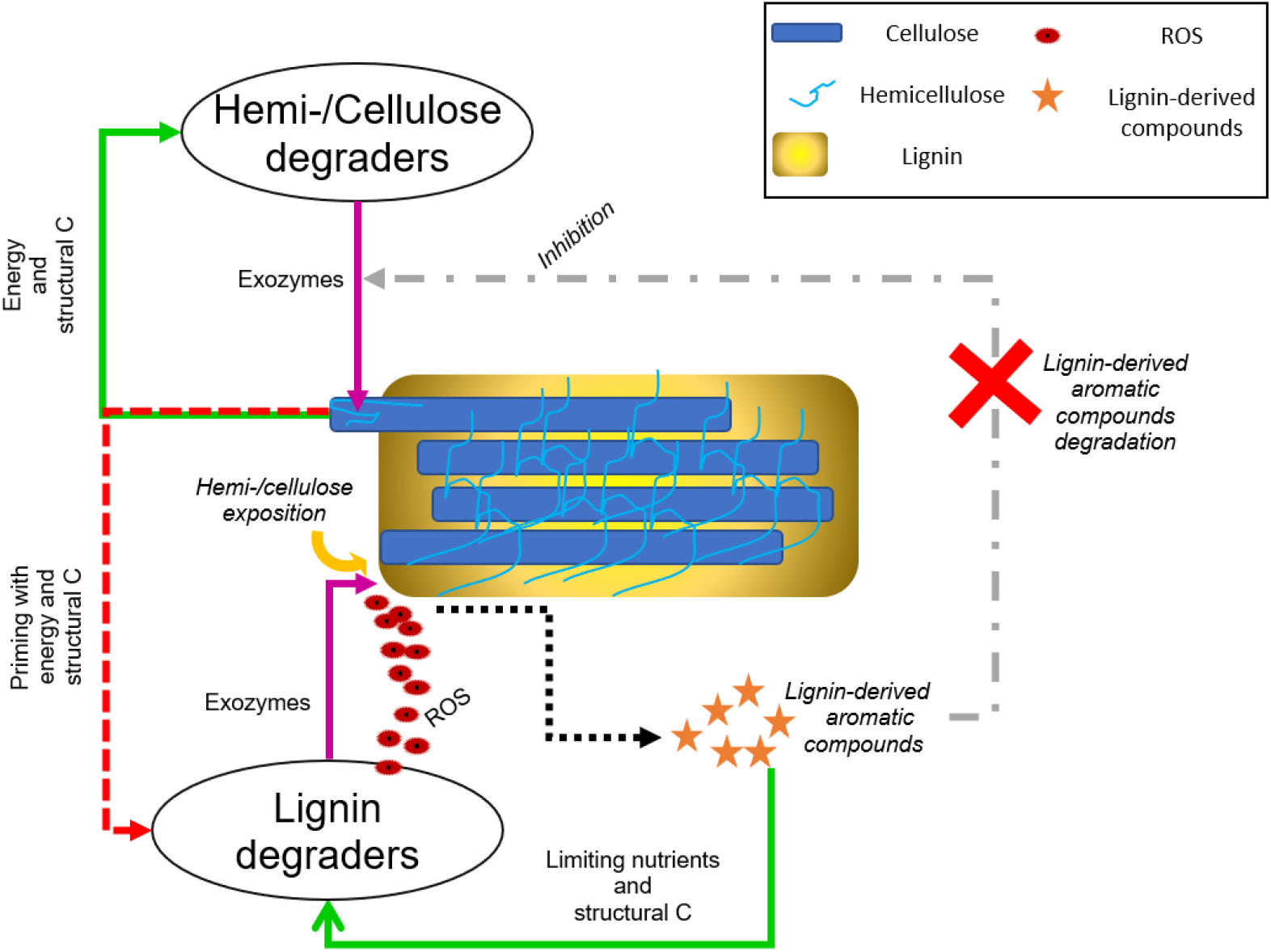
Priming effect model of microbial TeOM degradation in the Amazon River. The cellulolytic communities degrade hemi-/cellulose through secretion of glucosyl hydrolases (mainly GH3/GH10) which releases sugars to the environment. These sugars can promote growth in the cellulolytic and lignolytic communities, and during this process, the oxidative metabolism produces reactive oxygen species (ROS). ROS activate the exoenzymes (mainly through DYPs and laccases) secreted by the lignolytic community to oxidize lignin. After lignin oxidation, the hemi-/cellulose becomes exposed again, helping the cellulolytic communities to degrade it. During the previous process, several aromatic compounds are formed, which can potentially inhibit cellulolytic enzymes and microbial growth. However, these compounds are consumed by lignolytic microorganisms, reducing their concentration in the environment allowing decomposition to proceed. [Legend: green arrows – feedback; red dashed arrow – priming effect; black dashed arrow – products; magenta arrows – release of exoenzymes over a substrate; gray arrow – inhibition that cellulolytic organisms suffer from byproducts of lignin oxidation]

## MATERIALS AND METHODS

We analyzed 106 metagenomes^45–48^ from 30 stations distributed along the Amazon river basin, with an average coverage of 5.0×10^9^ base pairs per metagenome (standard deviation of 7.3×10^9^ bp / metagenome). The stations from Solimões River and lakes in the Amazon River course, located upstream from the city of Manaus, until the Amazon River’s plume in the Atlantic Ocean covered ~2,106 Km and were divided into 5 sections (**Fig. 1a, Supplementary Table 2**). These sections were: 1) *Upstream section* (upstream Manaus city); 2) *Downstream section* (placed between Manaus and the start of the Amazon River estuary. It includes the influx of particle-rich white waters from the Solimões River as well as the influx of humic waters from Negro River ^49, 50^), 3) *Estuary section* (part of the river that meets the Atlantic Ocean) and 4) *Plume section* (the area where the Ocean is influenced by the Amazon River inputs).

Samples were taken as previously indicated ^45–48^. Depending on the original study, particle-associated microbes were defined as those passing the filter of 300 μm mesh-size and being retained in the filter of 2 - 5 μm mesh-size. Free-living microbes were defined as those passing the filter of 2 - 5 μm mesh-size, being retained in the filter of 0.2 μm mesh-size. DNA was extracted from the filters as previously indicated^45–48^. Metagenomes were obtained from libraries prepared with either Nextera or TruSeq kits. Different Illumina sequencing platforms were used: Genome Analyzer IIx, HiSeq 2500 or MiSeq. Additional information is provided in **Supplementary Table 2**.

### Metagenome processing

Illumina adapters and poor quality bases were removed from metagenomes using Cutadapt^51^. Only reads longer than 80 bp, containing bases with Q ≥ 24, were kept. The quality of the reads was checked with FASTQC^52^. Reads from metagenomes belonging to the same station were assembled together using MEGAHIT (v1.0)^53^, with the meta-large presets. Only contigs > 1 Kbp were considered, as recommended by previous work^54^. Assembly quality was assessed with QUAST^55^.

### Analysis of k-mer diversity over different environments

A k-mer diversity analysis was used to compare the genetic information in the Amazon River microbiome against that in other microbiomes from Amazon forest soil or temperate rivers (**Supplementary Table 3**). Specifically, the Amazon River metagenomes (106) were compared against 37 metagenomes from the Mississippi River^56^, 91 metagenomes from three watersheds in Canada^57^, and 7 metagenomes from the Amazon forest soil^58^. The rationale to include soil metagenomes was to check whether genomic information in the river could derive from soils. K-mer comparisons were run with SIMKA (version 1.4)^59^ normalizing by sample size. Low complexity reads and k-mers (Shannon index < 1.5) were discarded before SIMKA analyses. The resulting Jaccard’s distance matrix was used to generate a non-metric multidimensional scaling (NMDS) analysis. Permutation tests were used to check the homogeneity of beta-dispersion in the groups, and permutational multivariate analysis of variance (PERMANOVA/ANOSIM) was used to test the groups’ difference. Both analyzes were performed using the R package Vegan^60^.

### Amazon River basin Microbial non-redundant Gene Catalogue (AMnrGC)

Genes were predicted using Prodigal (version 2.6.3)^61^. Only open reading frames (ORFs) predicted as complete (i.e. accepting alternative initiation codons, and longer than 150 bp) were considered in downstream analyses. Gene sequences were clustered into a non-redundant gene catalogue using CD-HIT-EST (version 4.6)^62, 63^ at 95% of nucleotide identity and 90% of overlap of the shorter gene^5^. Representative gene-sequences were used in downstream analyses. GC content per gene was inferred via Infoseq, EMBOSS package (version 6.6.0.0)^64^.

### Gene abundance estimation

The quality-trimmed sequencing reads were mapped to our non-redundant gene catalogue using BWA (version 0.7.12-r1039)^65^ and SamTools (version 1.3.1)^66^. Gene abundances were estimated using the software eXpress (version 1.5.1)^67^, with no bias correction, as the equivalent to transcripts per million (TPM) [Note that even though we use the common acronym TPM for simplicity, we have always used reads, no transcripts]. We used a TPM ≥ 1.00 for a gene to be present in a sample, and an average abundance higher than zero (μ_TPM_ > 0.0) for a gene to be present in a river section or water type (freshwater, brackish water or the mix of them in the plume).

### Functional annotation

Representative genes (and their predicted amino acid sequences) were annotated by searching them against KEGG (Release 2015-10-12)^68^, COG (Release 2014)^69^, CAMERA Prokaryotic Proteins Database (Release 2014)^70^ and UniProtKB (Release 2016-08)^71^ via the Blastp algorithm implemented in Diamond (v.0.9.22)^72^, with a query coverage ≥ 50%, identity ≥ 45%, e-value ≤ 1e-5 and score ≥ 50. KO-pathway mapping was performed using KEGG mapper^73^. HMMSearch (version 3.1b1)^74^ was used to search proteins against dbCAN (version 5)^75^, PFAM (version 30)^76^ and eggNOG (version 4.5)^77^ databases, using an e-value ≤ 1e-5, and posterior probability of aligned residues ≥ 0.9, and no domain overlapping. Accumulation curves were obtained using random progressive nested comparisons with 100 pseudo-replicates for genes and PFAM predictions.

### Gene taxonomic assignment

Gene-taxonomy was assigned considering the best hits (score, e-value and identity; see above) using KEGG (Release 2015-10-12)^68^, UniProtKB (Release 2016-08)^71^ and CAMERA Prokaryotic Proteins Database (Release 2014)^70^. Taxonomic last common ancestors (LCA) were determined from TaxIDs (NCBI) associated to UniRef100 and KO entries. Information from the CAMERA database was also used to retrieve taxonomy (NCBI TaxID). Proteins were annotated as ‘unassigned’ if their taxonomic signatures were mixed, containing representatives from several domains of life, or if they only had the function assigned without taxonomic information. Reference sequences with hits to poorly annotated sequences from other metagenomes were referred as “Metagenomic”.

### TeOM degradation machinery

To investigate the TeOM degradation, we grouped samples by river section and assessed their gene contents. These genes were then searched against reference sequences and proteins families characterized as TeOM degrading functions, shown in **Supplementary Table 4**.

Lignin degradation starts with extracellular polymer oxidation followed by internalization and metabolism of the produced monomers or dimers by bacteria. Protein families related to lignin oxidation (PF05870, PF07250, PF11895, PF04261 and PF02578) were searched among PFAM-annotated genes. The genes related to the metabolism of lignin-derived aromatic compounds were annotated with Diamond (Blastp search mode; v.0.9.22)^72^, with query coverage ≥ 50%, protein identity ≥ 40% and e-value ≤ 1e-5 as recommended by Kamimura et al.^37^, using their dataset as reference.

Cellulose and hemicellulose degradation involves glycosyl hydrolases (GH). The most common cellulolytic protein families (GH1, GH3, GH5, GH6, GH8, GH9, GH12, GH45, GH48, GH51 and GH74)^78^ and cellulose-binding motifs (CBM1, CBM2, CBM3, CBM6, CBM8, CBM30 and CBM44)^78, 79^ were searched in PFAM annotations. In addition, the most common hemicellulolytic families (GH2, GH10, GH11, GH16, GH26, GH30, GH31, GH39, GH42, GH43 and GH53)^79^ were searched in the PFAM database. Lytic polysaccharide monooxygenases (LPMO)^79^ were also identified using PFAM to investigate the simultaneous deconstruction of cellulose and hemicellulose.

During the degradation of refractory and labile material by exoenzymes, microbes produce a complex mix of particulate and dissolved organic carbon. The use of this mix is mediated by a vast diversity of transporter systems^38^. The typical transporters associated to lignin degradation (MFS transporter, AAHS family, ABC transporters, MHS family, ITS superfamily and TRAP transporter) were searched with Diamond (v.0.9.22)^72^, using query coverage ≥ 50%, protein identity ≥ 40% and e-value ≤ 1e-5 and a reference dataset previously compiled^37^.

Lignin degradation ends in the production of 4-carboxy-4hydroxy-2-oxoadipate, which is converted into pyruvate or oxaloacetate, both substrates of the tricarboxylic acid cycle (TCA)^37^, similarly to the fate of hemi-/cellulose degradation byproducts. Recently, several substrate binding proteins (TctC) belonging to the tripartite tricarboxylate transporter (TTT) system were related to the transporting of TeOM degradation byproducts, like adipate^39^ and terephtalate^40^. To investigate the metabolism of these compounds, and the possible link between the TTT system and lignin/cellulose degradation, the protein families TctA (PF01970), TctB (PF07331) and TctC (PF03401) were searched in PFAM.

The genes found using the above mentioned strategy were submitted to PSORT v.3.0^80^, to determine the protein subcellular localization (cytoplasm, secreted to the outside, inner membrane, periplasm, or outer membrane). We carried out predictions in the three possible taxa (Gram negative, Gram positive and Archaea), and the best score was used to determine the subcellular localization. Genes assigned to an “unknown” location, as well as those with a wrong assignment were eliminated (for example, genes known to work in extracellular space that were assigned to the cytoplasmic membrane).

The total amount of TeOM degradation genes found per function (lignin oxidation, transport, hemi-/cellulose degradation and lignin-derived aromatic compounds metabolism) in each section of the river, were normalized by the maximum gene counts per metagenome. Subsequently, correlograms were produced using Pearson’s correlation coefficients with the R packages Corrplot^81^ and RColorBrewer^82^. The linear geographic distance of each metagenome to the Amazon River source (Mantaro River, Peru, 10° 43′ 55″ S / 76° 38′ 52″ W), was also used in this analysis to infer changes in gene counts along the Amazon River course. The distance was calculated with the R package Fields^83^.

### Data availability

Metagenomes used to construct the Amazon River gene catalogue (AMnrGC) are publicly available (See **Supplementary Table 2**; SRA projects: SRP044326, PRJEB25171 and SRP039390). The AMnrGC, as well as, the annotation files are available in: 10.5281/zenodo.1484503. Metagenomes used in the k-mer diversity comparison are detailed in **Supplementary Table 3** (SRA projects: Amazon forest [PRJNA336764, PRJNA336766, PRJNA337825, PRJNA336700, PRJNA336765], Mississippi River [SRP018728] and Canada watersheds [PRJNA287840]).

## Acknowledgements

C.D.S.J. was supported by a PhD scholarship from Conselho Nacional de Desenvolvimento Científico e Tecnológico, Brazil (CNPq #141112/2016-6). F.H.S. and H.S work was supported by Research Productivity grants from CNPq (Process # 311746/2017-9 and #309514/2017-7, respectively). R.L. was supported by a Ramón y Cajal fellowship (RYC-2013-12554, MINECO, Spain). This work was supported by Petróleo Brasileiro S.A. (Petrobras), as part of a research agreement (#0050.0081178.13.9) with the Federal University of São Carlos, SP, Brazil, within the context of the Geochemistry Thematic Network. Additonally, this work was supported by the projects INTERACTOMICS (CTM2015-69936-P, MINECO, Spain) and MicroEcoSystems (240904, RCN, Norway) to RL and Fundação de Amparo à Pesquisa do Estado de São Paulo – FAPESP (Process #2014/14139-3) to HS. This study was financed in part by the Coordenação de Aperfeiçoamento de Pessoal de Nível Superior - Brasil (CAPES) - Finance Code 001 (CAPES #88881.131637/2016-01). Bioinformatics analyses were performed at the MARBITS platform of the Institut de Ciències del Mar (ICM; http://marbits.icm.csic.es) as well as in MareNostrum (Barcelona Supercomputing Center) via grants obtained from the Spanish Network of Supercomputing (RES) to RL. We thank Pablo Sánchez for his orientation with bioinformatics analyses and support. We also thank the EMM group (https://emm.icm.csic.es) at the ICM-CSIC for all the support and cordiality during the development of part of this work.

## Contributions

CDSJ, FHS & RL designed the study. CDSJ compiled and curated the data and performed bioinformatic analysis. CDSJ, FHS, HS & RL interpreted the results. FHS, RL, FPM and HS supervised and administered the project, providing funding. The original draft was written by CDSJ. All co-authors contributed substantially to manuscript revisions.

## SUPPLEMENTARY TABLES

**Supplementary Table 1. Co-assembly groups used to build the *Amazon River basin Microbial non-redundant Genes Catalogue* (AMnrGC).**

**Supplementary Table 2. Metagenomes used to build the *Amazon river basin Microbial non-redundant Genes Catalogue* (AMnrGC).** Description of the 106 metagenomes used in this study. The Amazon River basin region shows the group that a sample belongs according to its geographical location. Other features were obtained from the original publications and SRA. “N.A.” stands for not available.

**Supplementary Table 3. Metagenomes used for K-mer diversity assessment.**

**Supplementary Table 4. Reference proteins and Protein Families used in TeOM degradation functional searches.** PFAMs related to lignin oxidation, cellulose and hemicellulose degradation used to detect and annotate orthologous in the AMnrGC^37^.

