## Supplementary Figures for "Uncovering the gene machinery of the Amazon River microbiome to degrade rainforest organic matter"

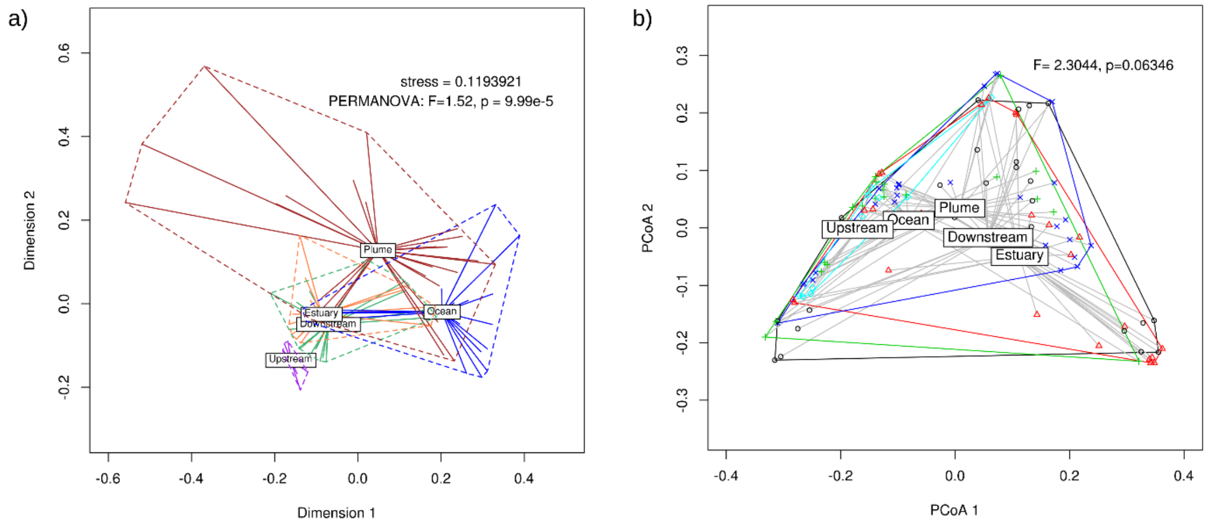


**Supplementary Figure 1. Metagenomic composition of the five studied sections of the Amazon River microbiome.** Ordination of metagenomes composing the different river sections based on the Jaccard distances calculated from the presence-absence of K-mers in each sample. NMDS groups were statistically different [PERMANOVA, F=1.52, p-value = 9.99e-5] (a), while the composition inside groups was homogeneous [β-dispersion; PERMUTEST, F=2.30, p-value = 0.06] (b).

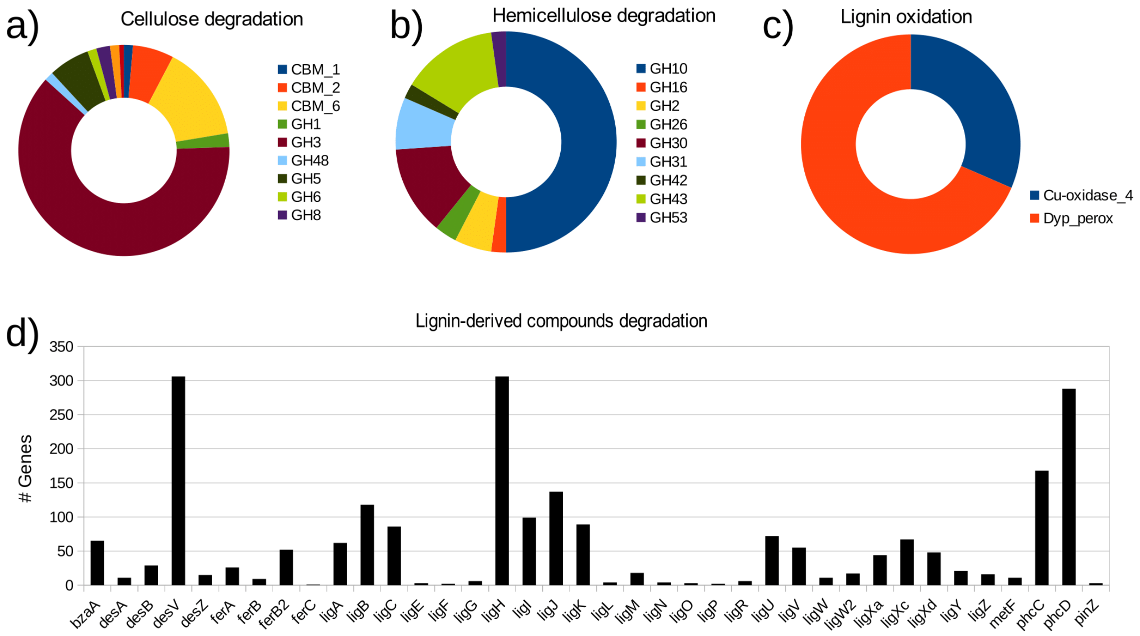


**Supplementary Figure 2. Main enzyme families involved in TeOM degradation in the AMnrGC.** a) Cellulose and b) hemi-cellulose degradation, c) lignin oxidation, and d) lignin-derived compounds degradation shown as number of detected genes per protein family. Abbreviations indicate protein families.
